# Translating outcomes from the clinical setting to preclinical models: chronic pain and functionality in chronic musculoskeletal pain

**DOI:** 10.1101/2021.10.27.466137

**Authors:** Melissa E. Lenert, Rachelle Gomez, Brandon T. Lane, Dana L. Dailey, Carol G.T. Vance, Barbara A. Rakel, Leslie J. Crofford, Kathleen A. Sluka, Ericka N. Merriwether, Michael D. Burton

## Abstract

Fibromyalgia (FM) is a chronic pain disorder characterized by chronic widespread musculoskeletal pain (CWP), tenderness, and fatigue, which interferes with daily functioning and quality of life. In clinical studies, this symptomology is assessed, while preclinical models of CWP are limited to nociceptive assays. The aim of the study was to investigate the human-to-model translatability of clinical behavioral assessments for pain and muscle function in a preclinical model of CWP. We assessed correlations between pain behaviors and muscle function in a preclinical model of CWP and in women with fibromyalgia to examine whether similar relationships between outcomes existed in both settings, for usability of clinical assays in model systems. For preclinical measures, the acidic saline model of FM which induces widespread muscle pain, was used in adult female mice. Two gastrocnemius injections of acidic or physiological pH saline were given following baseline measures, five days apart. An array of adapted pain measures and functional assays were assessed for three weeks. For clinical measures, pain and functional assays were assessed in adult women with FM. For both preclinical and clinical outcomes, movement-evoked pain (MEP) was associated with mechanical pain sensitivity. Mechanical sensitivity was correlated to shifts in weight-bearing preclinically and was predictive of functionality in patients. Preclinically, it is imperative to expand how the field assesses pain behaviors when studying multi-symptom disorders like FM. Targeted pain assessments to match those performed clinically is an important aspect of improving preclinical to clinical translatability of animal models.

**Summary:** Preclinical assessments of chronic musculoskeletal pain recapitulate several outcome measures for clinical assessment of patients with FM, particularly prolonged resting pain, and MEP.

## 1. Introduction

Within the chronic pain field, there is often a disconnect between successful preclinical therapeutic strategies and those that are successful in clinical trials (1, 2). One of the contributing factors to this disconnect may be how pain is assessed in animal models compared to patient outcomes measured in clinical trials (3, 4). Behavioral assays in animal models primarily focus on evoked pain measures whereas clinical trials primarily focus on the assessment of self-reported spontaneous pain and functional outcomes (5, 6). To better identify potential therapeutics, it is essential that preclinical assays are designed to improve translatability to clinical phases of testing and implementation.

Fibromyalgia (FM) is a chronic musculoskeletal pain disorder classified by widespread pain, fatigue, and cognitive symptoms that has a higher prevalence in women than men (7). Preclinical studies of FM have greatly advanced over the years with the development of multiple animal models that mimic widespread pain (8). The acidic saline model was developed in 2001 as a non-inflammatory model of chronic widespread musculoskeletal pain (CWP). It consists of two unilateral injections of low pH saline into the calf muscle to produce widespread, long-lasting hyperalgesia without local tissue damage and inflammation (9). As FM is a multi-symptom disorder, there are several aspects of quality of life that are impaired; however, the focus of preclinical studies is primarily limited to studies of evoked pain (7, 10–12). These assays do not capture the full spectrum of symptoms experienced by patients with FM. Improving the translatability of assessments of pain and physical function used in the preclinical phase by utilizing assays that measure outcomes like those measured in clinical studies is imperative for improving success rates in clinical trials and maximizing patient outcomes. In addition to tests for nociception in rodents, the use of functional assays provides a more comprehensive assessment of the effects of chronic pain on function (5). With CWP, muscle function is often impaired and leads to worsening of pain and fatigue in patients (13, 14). Pain and fatigue are also worsened during physical activity (14). Many pain interventions for patients with FM feature multimodal non-pharmacological treatment strategies that primarily target myofascial pain and fatigue (15); However, the role of muscle performance and movement-evoked pain (MEP) in animal studies of FM have not been adequately characterized and limited to muscle fatigue (16). Targeted alleviation of MEP and fatigue may improve the efficacy of exercise-based treatments, which are effective in both clinical and preclinical studies (17–19).

Thus, the purpose of this study was to characterize spontaneous pain, MEP, and muscle performance in a preclinical model of CWP, and validate behavioral assays using clinical data. Using the acidic saline model, this study shows that CWP induces prolonged spontaneous pain behaviors, MEP, and reduced muscle strength in female mice. We examined correlative relationships between these assays and found that spontaneous pain and MEP are strongly related. To understand the translational value of these assays, we then examined relationships among assays used in clinical assessment of women with FM, between pain and patient-reported outcomes.

## 2. Materials and Methods

### 2.1 Animals

Female C57BL/6J mice (2-5 months old, 18-25 g) were used for the current study. Females were chosen to match the clinical cohort which used females with FM. Mice were purchased from Jackson Laboratory (stock no. 000664) and bred in-house at UT-Dallas. Mice were group housed (4-5 per cage) in polypropylene cages and maintained at a room temperature of 21 ± 2°C under a 12-hour (lights on from 6AM to 6PM) light-dark cycle with ad libitum access to water and rodent chow. Mice were handled daily for two weeks and habituated to the testing room prior to all experiments. All procedures were in accordance with the National Institutes of Health Guidelines for the Care and Use of Laboratory Animals and approved by the University of Texas at Dallas Institutional Animal Care and Use Committee.

### 2.2 Induction of Chronic Widespread Pain (CWP)

Two gastrocnemius injections of acidic saline (0.9%, pH 4.0 ± 0.1) were used to induce chronic bilateral sensitivity, as described previously (9, 11). Briefly, mice were anesthetized with isoflurane (5% induction, 2.5% maintenance) and the left gastrocnemius muscle was injected with 20µL of pH 4.0 (± 0.1) or pH 7.4 (± 0.1) saline on Days 0 (following baseline measures) and 5.

### 2.3 Behavior Testing

The day prior to baseline measures, animals were acclimated to the testing room for four hours. On testing days, animals were acclimated for approximately one hour in acrylic behavior boxes prior to testing. Behavior racks were cleaned with a 1:3 ratio of deodorant-free cleaner (Seventh Generation ^TM^, 22719BK-5) to eliminate odor cues. All behavioral tests were performed with the experimenter blinded to group. Unless otherwise stated, behavioral testing occurred between 1PM and 5PM. Timelines for behavioral tests can be found in Figure 1, as well as within each corresponding figure.

**Figure 1.**
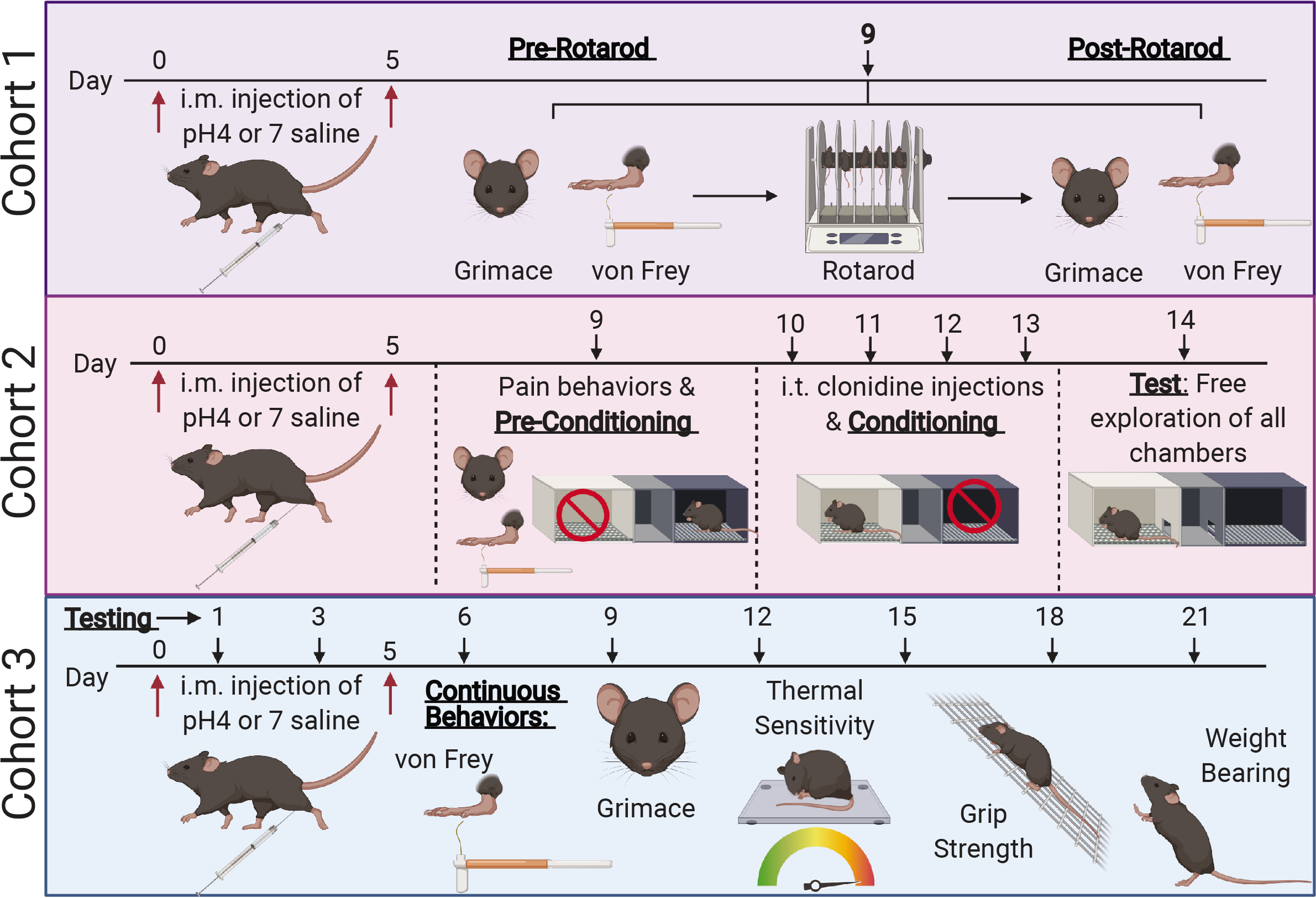
Representative timelines for preclinical behavioral assessments.

#### 2.3.1. Movement-evoked Pain (MEP)

Nine days following the first acidic saline injection, we administered the rotarod test as a means of short-term activity to induce MEP. MEP was assessed on day nine by measuring grimace scores and mechanical hypersensitivity (described in Sections 2.3.2 and 2.3.3, respectively) at baseline, before, and after forced activity on a rotarod apparatus (IITC Life Science, Series 8) (20). Mice were acclimated to the testing room for one hour prior to testing during the beginning of the dark phase of the 12-hour light/dark cycle, between 6 PM and 9 PM. Prior to baseline measures, mice were habituated to the stationary rotarod apparatus for five minutes each. On day nine after pain assessments (mechanical sensitivity of the paw and mean grimace score (MGS) pre-movement), mice were placed on the stationary rotarod and performed three back-to-back trials using the following settings: starting speed was 10 rotations per minute (RPM), 30 second ramp speed, and 45 RPM maximum speed. The time until each animal fell from the rod was recorded automatically by the apparatus for each trial. The rod is at a height such that mice are motivated to stay on the rotating rod but are unharmed when falling off it. Mice were immediately placed back into acrylic behavior boxes after the third trial. Ten minutes following activity, grimacing and mechanical hypersensitivity were assessed again (mechanical sensitivity of the paw and MGS post-movement). Percent change (MGS) was determined by calculating the percent change in grimace after the rotarod (MGS post-movement) to grimace before the rotarod (MGS pre-movement).

#### 2.3.2 Grimace

Spontaneous pain was assessed using the mouse grimace scale, in which the experimenter rates aspects of facial expression (orbital, cheeks, nose, whiskers, and ears) on a scale of 0-2, which are averaged to give the mean grimace score (MGS) (21). A score of “0” indicates facial grimacing is not present, “1” indicates moderate facial grimacing, and “2” indicates obvious facial grimacing. Animals were acclimated in acrylic behavior boxes for one hour prior to assessment of facial grimacing. Grimace scores were assessed at baseline (prior to any other tests performed), and every three days following for three weeks.

#### 2.3.3 von Frey Testing

Mechanical hypersensitivity was tested using the von Frey assay, described previously (11, 22). Animals were habituated to acrylic behavior boxes for a minimum of one hour prior to baseline measures. Paw withdrawal thresholds were assessed using calibrated von Frey hair filaments using the up-down method (22). Filaments with logarithmically incremental stiffness (2.83, 3.22, 3.61, 3.84, 4.08, 4.17 converted to 0.07, 0.16, 0.4, 0.6, 1, 1.4 g, respectively) were applied to the plantar surface of the hind paw. A 2.0 g cut off was applied to avoid tissue damage or unintentional agitation. A positive response was noted by paw withdrawal, licking, or shaking of the paw. Testing of the ipsilateral and contralateral paws occurred separately.

#### 2.3.4 Conditioned Place Preference (CPP)

We assessed persistent pain via analgesia-induced place preference using clonidine, a sedative and antihypertensive that has been used to treat a variety of pain conditions (23). CPP was performed following a previously published protocol (24). The CPP apparatus contained three chambers: one black, one white, and a striped connecting chamber. All CPP experiments were conducted between 6PM and 11PM. CPP was conducted in a dark room with a single low lumens light source (Maia 69E120-WH). The two end chambers were differentiated by scent (differently scented chapstick applied to the ceiling) and floor texture. The middle chamber was brightly lit to discourage mice from spending time in this chamber. Before pre-conditioning, animals were tested for mechanical sensitivity and facial grimacing (Day 9). During preconditioning, mice freely explored the middle and black chambers for 10 minutes. Conditioning occurred on Days 10-13. On conditioning days, mice were deeply anesthetized with isoflurane (5% induction, 2.5% maintenance) and injected intrathecally with clonidine (10µg/5µL) or vehicle (1X PBS, 5µL) and confined to the white chamber for 30 minutes. Successful analgesia via clonidine will elicit a preference for the white chamber in mice experiencing chronic pain, but not in pain-free controls. Additionally, mice that do not experience analgesia (vehicle injected groups) will not have any preference for the clonidine-paired chamber. There was approximately one hour between injections and conditioning. On testing day (Day 14), each mouse was placed in the middle chamber with unrestricted access to all chambers and recorded for ten minutes. The CPP apparatus was cleaned with 70% ethanol between mice to prevent odor cues.

Data are presented as a proportion of time spent in the drug-paired chamber over time spent in the unpaired chamber. Percent change analysis is presented as the percent change in chamber preference compared to the control group (pH7, vehicle) and calculated as Percent Change = ((Chamber Preference/Control Group Average) *100)-100. Values greater than zero indicate increased preference for the drug-paired chamber compared to control. Locomotor behavior, measured as the number of crossings between chambers, was used as an exclusionary criterion via outlier analysis. All data was scored by experimenters blinded to condition.

#### 2.3.5 Thermal Sensitivity

To assess changes in heat sensitivity, mice were tested for withdrawal latency to a noxious temperature. Mice were placed in an acrylic box with a temperature-controlled metal plate that was heated to 52°C (IITC). The latency (in seconds) for the mouse to withdraw its hind paw was recorded. Response latency was assessed at baseline, and every three days following the first injection for three weeks.

#### 2.3.6 Muscle Strength and Weight Bearing (incapacitance)

Hind and forelimb muscle performance were assessed using the grip strength assay and the incapacitance assay. The grip strength assay was performed as previously described (25, 26). Briefly, mice were suspended by the tail, allowed to grasp a wire mesh, and gently pulled backward. The maximum force (g) generated by the grip strength meter (IITC) was recorded. The incapacitance assay measures weight bearing in rodents. Mice were placed in the incapacitance meter (IITC) with one hind paw on each platform. The average weight (g) on each platform was recorded for a testing period of ten seconds, giving a measurement of weight distribution in a resting state, represented as a ratio of weight on the ipsilateral paw over the contralateral paw (27). Values of “1” indicate equal preference, greater than “1” indicate ipsilateral preference, and less than “1” indicate contralateral preference.

#### 2.3.5 Muscle coordination and endurance

The rotarod test was used to measure endurance and coordination following acidic saline injection (20). Mice were tested at baseline and at nine and 21 days following the first saline injection. All testing occurred during the dark cycle (6PM-9PM). Prior to baseline measures, mice were habituated to the stationary rotarod apparatus for five minutes each. Tests were run using the settings described in section 2.3.1. Each mouse performed three back-to-back trials for each time point measured. The time until each animal fell was recorded for each trial.

### 2.4 Human Studies

#### 2.4.1. Study Design

The human subjects’ study is a secondary analysis of data at baseline from participants enrolled in a two-site randomized, controlled clinical trial of the efficacy of transcutaneous electrical nerve stimulation (TENS) on MEP in women with fibromyalgia (FM). (Fibromyalgia Activity Study with TENS, FAST; ClinicalTrials.gov identifier NCT01888640; registered on June 28, 2013). The study was approved by the institutional review boards of both study sites. A detailed study protocol has been previously published (28).

#### 2.4.2. Study Participants

Briefly, women with FM were recruited from the University of Iowa Hospitals and Clinics and the Vanderbilt University Medical Center and their surrounding communities. Study inclusion criteria were as follows: (1) English-speaking, (2) adults between 18 and 70 years old, (3) diagnosed with FM based on the 1990 and 2011 American College of Rheumatology criteria (the study was completed prior to the updated 2016 FM diagnostic criteria), and (4) reported an average pain rating of 4 or higher on the Numeric Rating Scale (NRS) over the past 7 days. Participants with FM were excluded if they reported an average pain intensity of less than 4 out of 10 over the past 7 days, reported previous TENS use within the past 5 years, had neuropathy or an autoimmune disorder, or were pregnant. Additional detailed information about study participants is previously published (11, 17) . Data from 289 women were included in the analyses.

### 2.5. Data Collection and Analysis

All tests and measures were performed by trained research personnel. Demographic data and baseline tests and measures were collected during two visits. During Visit 1, participant demographics were collected using an online questionnaire. At Visit 2, participants completed a battery of surveys and pain tests and measures. Data from all surveys and dynamic laboratory pain tests were recorded using the Research Electronic Data Capture (REDCap) system.

#### 2.5.1 Pain Intensity and Pain Interference

Pain intensity at rest (spontaneous pain) and pain interference were measured using the NRS and the Brief Pain Inventory. Anchors for the NRS are 0 = “no pain” through 10 = “worst pain imaginable.” Pain severity and pain interference were assessed using the Brief Pain Inventory (BPI) Short Form. The BPI is a 15-item survey with two subscales (Severity, Interference) that query the magnitude and impact of pain. Pain interference was also assessed using a PROMIS Pain Interference Short-Form.

#### 2.5.2. Mechanical Pressure Pain Threshold (PPT)

Pressure mechanical pain threshold was quantified using a digital pressure algometer (Somedic AB, Farsta Sweden) to assess hyperalgesia. Mechanical pressure was applied to the cervical, lumbar, and right lower leg regions at a rate of 40 kPa per second. Lower PPTs are indicative of higher pain sensitivity, and thus, hyperalgesia (29).

#### 2.5.3. Conditioned Pain Modulation (CPM)

CPM was assessed to determine the function of descending inhibitory pain modulation which is analogous to a “pain inhibiting pain” phenomenon. Methods have been described previously (11). In brief, CPM was measured using mechanical PPT as the test stimulus before and after cold water immersion (conditioning stimulus). Mechanical PPTs were measured at the lumbar and right lower leg sites. Values for CPM were expressed as a ratio of PPT values at the lumbar and right lower leg before and after cold water immersion (post PPT/pre PPT values). A reduction or no change in PPT values after cold-water immersion (ratio > 1) indicates impaired descending pain inhibition (30).

#### 2.5.4. Five time Sit-to-Stand Test (5TSTS)

The Five-Times-Sit-to-Stand Test (FTSTS) measures the time (seconds) for participants to transition from a seated position to a fully standing position five times. A higher time indicated the individual required more time to go from the seated position to a stance five times (31, 32).

#### 2.5.5. Six-minute walk test (6MWT)

The 6MWT is an assessment of physical endurance. Participants were instructed to walk for 6 minutes over a 50-foot walkway and total distance in feet was measured. The 6MWT has been well validated in chronic pain populations (33, 34).

#### 2.5.6. Movement-Evoked Pain (MEP)

MEP was assessed using the NRS (0-10) during the Six-Minute Walk Test (6MWT) and the Five-Times-Sit-to-Stand Test (FTSTS) (17). To assess MEP during the 6MWT, participants were asked to rate their pain at the 5-minute mark using the NRS. To assess MEP during the 5TSTS, participants were asked to rate their pain immediately after completing the task.

### 2.6 Statistical Analysis

#### 2.6.1. Animal Data

Data are presented as mean ± SEM. Correlative relationships were determined using Pearson R values and statistical comparisons were made using GraphPad Prism 8.4 statistical software for all preclinical datasets. Statistical significance was set at p ≤ .05. Repeated measures two-way ANOVA with post hoc Sidak’s multiple comparisons was used to assess the effects of acidic saline injections on pain (MEP, mechanical sensitivity, facial grimacing, and thermal sensitivity) and functional assays (grip strength, weight bearing, and rotarod). Data from CPP experiments (chamber preference and percent change) were analyzed using Ordinary One-Way ANOVA with post hoc Tukey’s multiple comparisons.

Effect sizes for MEP, facial grimacing, mechanical and thermal sensitivity, grip strength, and weight-bearing assays were determined by calculating the cumulative difference between baseline and each time point (26, 35, 36). Group comparisons were performed using an unpaired two-tailed t-test or Ordinary One-Way ANOVA, with statistical significance set at p < 0.05.

#### 2.6.2. Human Subjects Data

Descriptive summary statistics for the demographic and patient-reported outcome measures were calculated. Data are presented in their original units, and normality was determined using a Kolmogrov-Smirnoff test. The associations between outcome measures were calculated using Spearman Rho correlation coefficients. All analyses were conducted using SPSS version 25. Statistical significance was set at p ≤ 0.05.

### 3. Results

#### 3.1 Movement-evoked facial grimacing in mice with chronic widespread pain

MEP experienced by patients with FM is often not treated effectively by current therapeutics and can be a major limitation in exercise-based therapeutics (17). We investigated whether MEP was recapitulated in the preclinical acidic saline model of CWP by investigating the effects of activity on facial grimacing and mechanical sensitivity. Nine days following the first acidic saline injection, we administered the rotarod test as a means of short-term activity to induce MEP. Facial grimacing and mechanical hypersensitivity were assessed before and after multiple trials on the rotarod apparatus (Figure 2A).

**Figure 2.**
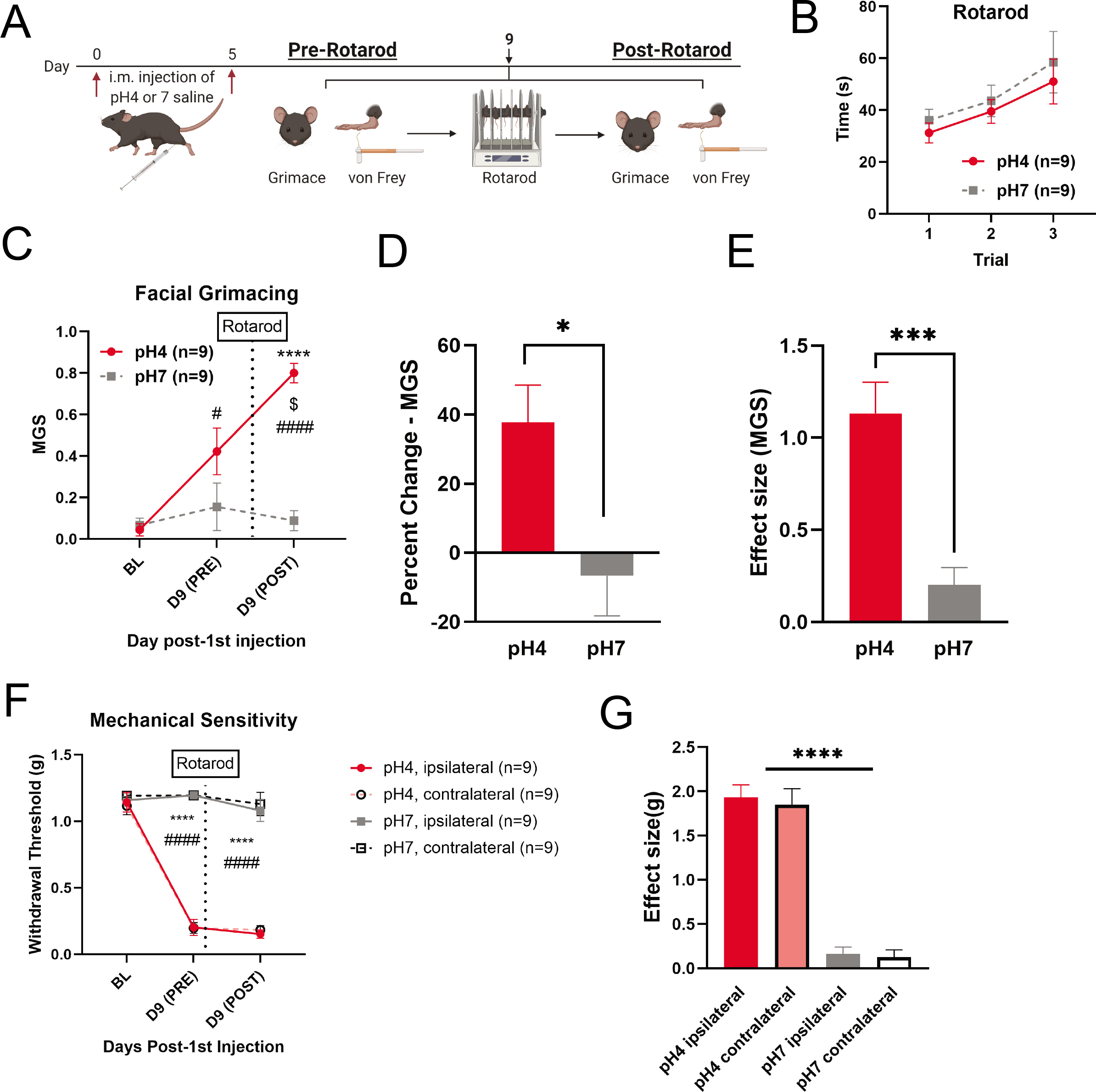
Movement-evoked pain in the acidic saline model of FM. (A) Timeline for behavioral assessments. Mice were given injections of pH4 or 7.4 saline into the gastrocnemius muscle on Days 0 and 5. On Day 9, mice were assessed for facial grimacing and mechanical sensitivity before and after multiple back-to-back trials on the rotarod apparatus. (B) There are no group differences in time spent on the rotarod. (C) Facial grimacing significantly increases after movement on the rotarod in acidic saline treated, but not control, mice (time x treatment: F (2,32) = 13.59, p < 0.0001, Repeated Measures Two- Way ANOVA, n=9/group). * pH4 vs pH7. # within pH4 group comparison to baseline. $ within pH4 group pre- vs post-rotarod. (D) Acidic saline treated mice develop bilateral sensitivity, but there is no change in mechanical sensitivity after movement on the rotarod (time x treatment: F (6, 64) = 29.02, p < 0.0001, Repeated Measures Two-Way ANOVA, n=9/group). *pH4 vs pH7 ipsilateral. # pH4 vs pH7 contralateral. (E) Percent change in facial grimacing pre- and post-rotarod. Acidic-saline treated mice have a significant increase in facial grimacing after movement compared to controls (t (16) = 2.814, p = 0.0125, unpaired t- test, two-tailed, n=9/group). (F) Effect size for facial grimacing. Acidic saline treated mice grimace significantly more across the entire experiment compared to controls (t (16) = 4.802, p = 0.0002, unpaired t-test, two-tailed, n=9/group). (G) Effect size for mechanical sensitivity. Acidic saline treated mice have significantly greater mechanical sensitivity compared to controls (F (3, 32) = 61.17, p < 0.0001, Ordinary one-way ANOVA, n=9/group). *pH4 vs pH7. All data are represented as mean ± SEM. *p<0.05, **p<0.01, ***p<0.001, ****p<0.0001. Graphics were created using BioRender.com.

Both groups showed the same level of improvement in time spent on the rotarod apparatus over the three trials (Figure 2B: trial: F (1.409, 19.72) = 6.317, p = 0.0133). There was a main effect of group on facial grimacing (Figure 2C) and mechanical sensitivity of the paw. Furthermore, there was an interaction effect of group and activity such that mice that received acidic saline injections showed a significant increase in facial grimacing after activity, whereas control mice showed no changes in facial grimacing (Figure 2C: time x treatment: F (2,32) = 13.59, p < 0.0001). There was also an interaction effect of group and activity such that mice that received the acidic saline injections showed a significant decrease in mechanical hypersensitivity in both paws compared with control mice (Figure 2D: time x treatment: F (6,64) = 29.02, p < 0.0001). However, post hoc comparisons showed no change in mechanical hypersensitivity of the paw after the rotarod test in either group. Furthermore, percent change analysis of facial grimacing (pre-movement vs post-movement) showed that mice that were treated with acidic saline showed a marked increase in facial grimacing after the rotarod test (Figure 2E: t (16) = 2.814, p = 0.0125). Analysis of effect size for MGS (cumulative difference from baseline) showed a robust increase in facial grimacing across the entire experiment (Figure 2F: t (16) = 4.802, p = 0.0002, unpaired t-test, two-tailed), whereas control mice had no changes in grimacing throughout the experiment. Comparisons of effect sizes for mechanical sensitivity of the paw showed group differences in mechanical sensitivity, but no differences between ipsilateral and contralateral sensitivity were observed for either group (Figure 2G: F (3,32) = 61.17, p < 0.0001, ANOVA), indicating that activity increases spontaneous pain in this model.

#### 3.3 Gastrocnemius injections of acidic saline induces prolonged spontaneous pain

Due to previous observations that the acidic saline model of CWP induces prolonged bilateral sensitivity (11), we investigated the presence of spontaneous pain using the CPP assay. CPP was used as a measure for spontaneous pain using analgesia as the conditioning stimulus. Place preference for the analgesic- paired chamber (clonidine) in mice with CWP, but not controls, would indicate that clonidine treatment alleviates the aversive qualities of CWP without having a rewarding effect in control mice. Mice that received acidic saline and clonidine (i.t., 10µg/5µL) exhibited preference for the clonidine-paired chamber, whereas the animals that received vehicle treatments did not (Figure 3B, F (3,24) = 5.893, p = 0.0037, ANOVA). When compared using percent change, mice that received acidic saline and clonidine spent double the amount of time in the drug-paired chamber compared to each control group (Figure 3C, (F (3, 23) = 3.748, p = 0.0251, ANOVA), indicating that the acidic saline model induces spontaneous pain that is successfully relieved by clonidine treatment.

**Figure 3.**
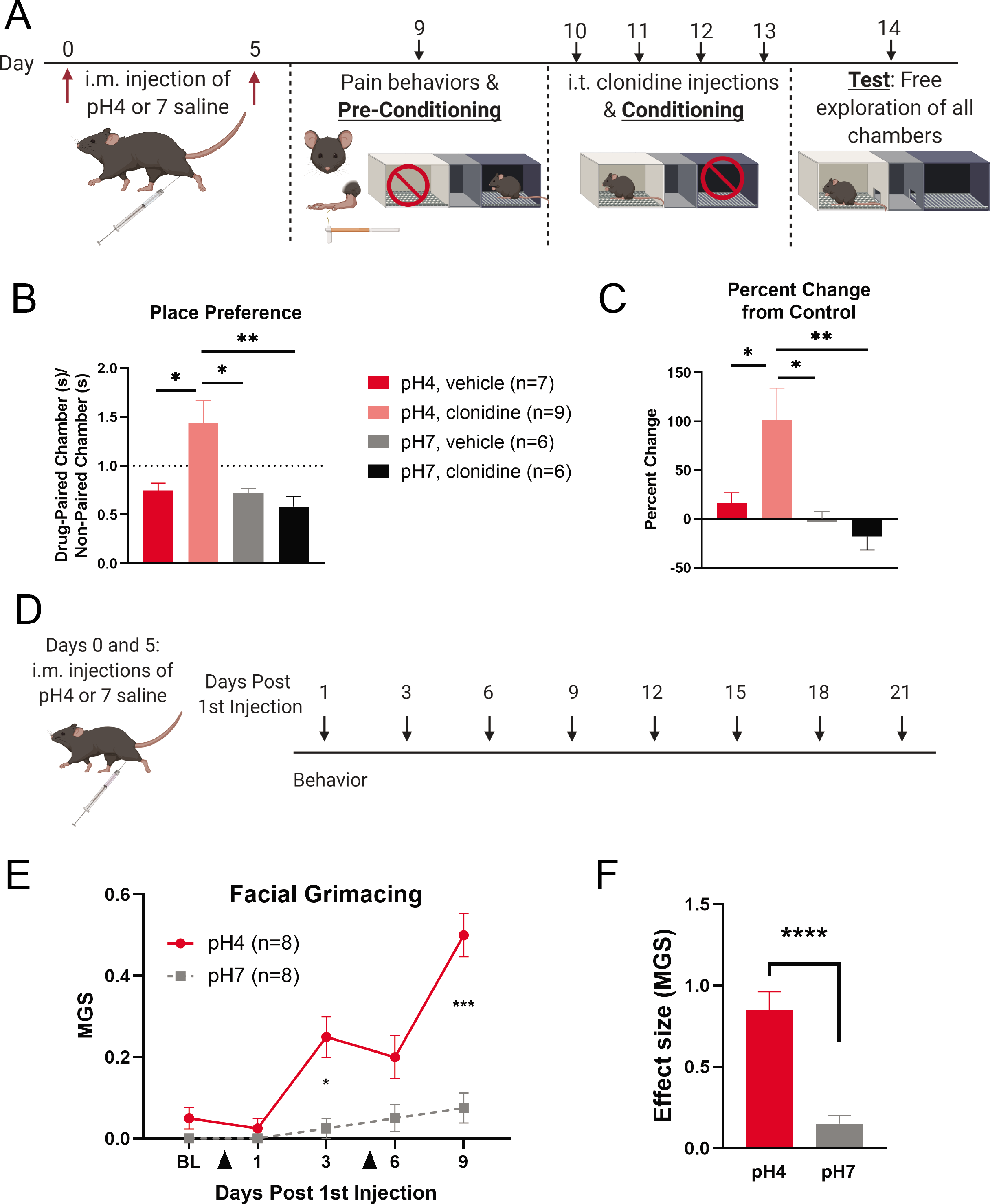
Spontaneous pain in the acidic saline model of FM. (A). Timeline for CPP experiments. Mice were given injections of pH4 or 7.4 saline into the gastrocnemius muscle on Days 0 and 5. Preconditioning occurred on Day 9 after pain behavior assessments. Conditioning via i.t. injections of clonidine (10ug/5uL) or vehicle (1X PBS, 5uL) occurred on Days 10-13. On Day 14, animals were allowed to freely explore the entire apparatus for 10 minutes and were video recorded. (B) Clonidine induces place preference in acidic saline treated mice, but not controls (F (3, 24) = 5.893, p = 0.0037, Ordinary One-Way ANOVA). (C) Percent change analysis of CPP score compared to the control group (pH7, vehicle) (F (3, 23) = 3.748, p = 0.0251, Ordinary One-Way ANOVA). (pH4, vehicle: n=7; pH4, clonidine: n=9; pH7, vehicle: n=6; pH7, clonidine: n=6). (D) Timeline for assessment of facial grimacing. Mice were given injections of pH4 or 7.4 saline into the gastrocnemius muscle on Days 0 and 5. Facial grimacing was assessed every three days. (E) Mice that received acidic saline injections have prolonged facial grimacing behavior compared to mice that received physiological saline injections (Time x Treatment: F (4, 56) = 9.915, p < 0.0001, Repeated Measures Two-Way ANOVA, n=8/group). Black arrowheads indicate saline injections. (F) Effect size analysis shows increased facial grimacing behavior over the measured time course (t (6) = 6.582, p = 0.0006, unpaired t-test, two-tailed, n=8/group). All data are represented as mean ± SEM. MGS = mean grimace score. *p<0.05, **p<0.01, ***p<0.001, ****p<0.0001.

We investigated the effect of CWP induced by acidic saline injections on facial grimacing behavior to assess spontaneous pain. Grimacing behavior is used to quantitatively assess changes in spontaneous ongoing pain over time without the need for using stimuli (such as mechanical stimuli) to evoke a response. Facial grimacing is typically short-lasting in mice and is robust during acute pain states (37). CWP induces prolonged facial grimacing that peaks at Day 9-post injection (Figure 3E, time x treatment: F (4, 56) = 9.915, p < 0.0001, ANOVA) and lasts for several weeks (Supplemental Figure 1A, time x treatment: F (8, 48) = 3.318, p = 0.0043, ANOVA). The effect size for grimacing behavior is significantly increased compared to controls through Day 9 post-injection (Figure 3F, t (14) = 5.715, p < 0.0001, unpaired t-test, two-tailed) and Day 21 post-injection (Supplemental Figure 1B, t (6) = 7.667, p = 0.0003, unpaired t-test, two-tailed).

#### 3.3 Gastrocnemius injection of acidic saline induces prolonged reduction in thermal sensitivity and muscle function

To investigate the effects of CWP on additional measures of evoked pain, we measured the bilateral response latency to a noxious temperature. Acidic saline treatment reduces response latency (Figure 4B, treatment: F (1,14) = 7.745, p = 0.0147, ANOVA) through three weeks of testing (Supplemental Figure 1C, treatment: F (1,14) = 11.72, p = 0.0041, ANOVA), with analysis showing an interaction between time and saline treatment over the three-week period (Supplemental Figure 1C, time x treatment: F (8,80) = 4.624, p = 0.0001, ANOVA). The effect size analysis for this assay was not significant (Figure 4C).

**Figure 4.**
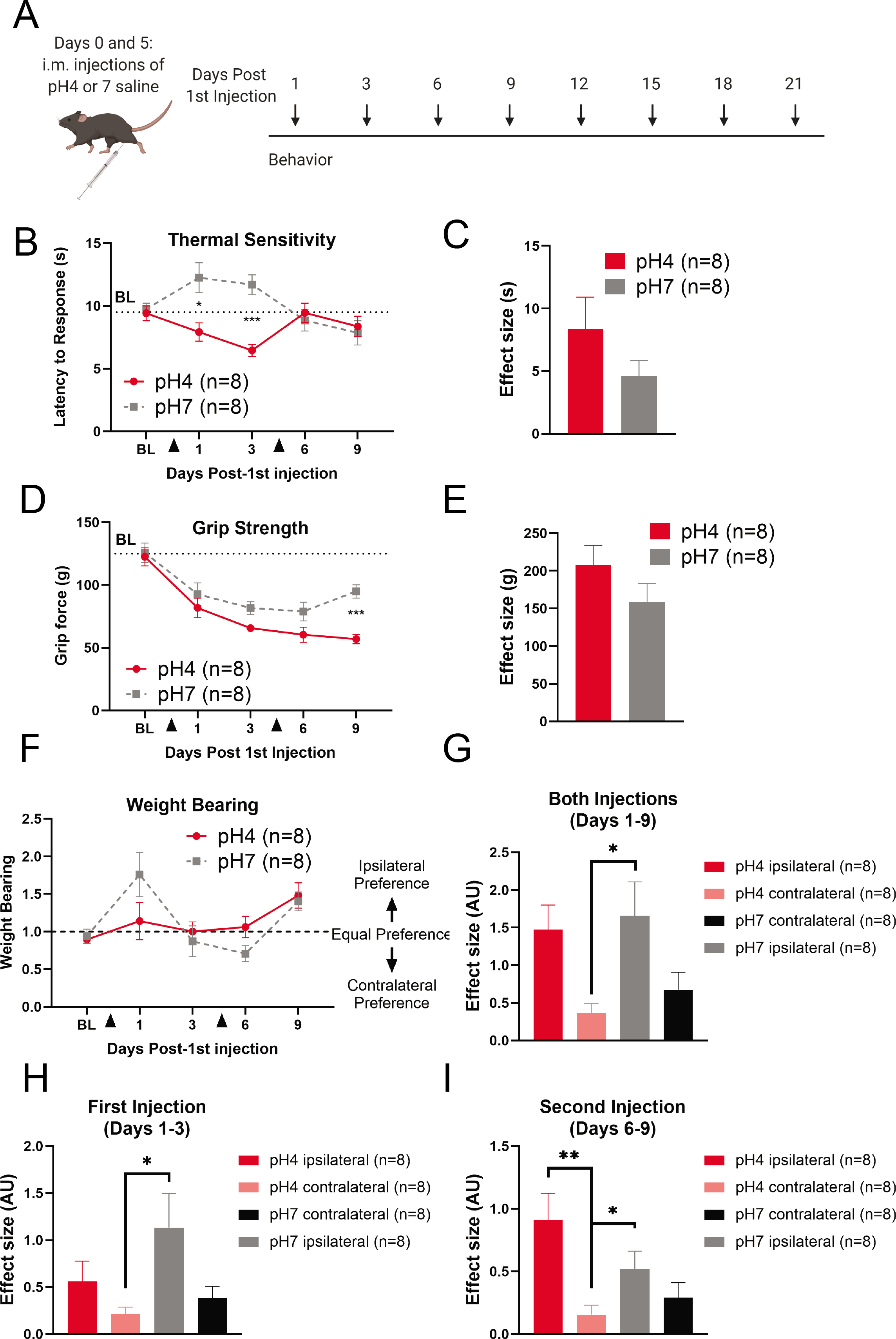
Assessment of thermal sensitivity and muscle strength in experimental model of chronic musculoskeletal pain. (A) Timeline for behavioral experiments. Mice were given injections of pH4 or 7.4 saline into the gastrocnemius muscle on Days 0 and 5. Behaviors were assessed every three days. (B) Mice that received gastrocnemius injections of acidic saline show an increase in sensitivity to noxious temperature (F (1,14) = 7.745, p = 0.0147, Repeated Measures Two-Way ANOVA, n=8/group). (C) Effect size analysis for thermal sensitivity. (D) Mice that received gastrocnemius injections of acidic saline show decreased grip strength (treatment: F (1,14) = 14.44, p = 0.0020, Repeated Measures Two-Way ANOVA, n=8/group). (E) Effect size analysis for grip strength. (F). Mice that received gastrocnemius injections of acidic saline show shifts in weight bearing at rest (time: F (2.664, 37.31) = 6.053, p = 0.0026, Repeated Measures Two-Way ANOVA, n=8/group). (G) Effect size analysis of total shifts in weight bearing (F (3, 28) = 4.108, p = 0.0170, Ordinary One-Way ANOVA, n=8/group). (H) Effect size analysis of shifts in weight bearing after the first injection (F (3, 28) = 3.228, p = 0.0374, Ordinary One-Way ANOVA, n=8/group). (I) Effect size analysis of shifts in weight bearing after the second injection (F (3, 28) = 5.137, p = 0.0059, Ordinary One-Way ANOVA, n=8/group). Graphics were created using BioRender.com. Black arrowheads indicate saline injections. All data are represented as mean ± SEM. *p<0.05, **p<0.01, ***p<0.001.

To investigate the effects of CWP on muscle function, we assessed the efficacy of assays designed to test different aspects of muscle performance. The grip strength test, which measures the force with which mice use their limbs to grip onto a metal grid, showed a significant effect of acidic saline on grip strength (Figure 4D, treatment: F (1,14) = 14.44, p = 0.0020, ANOVA) but not a significant interaction (time x treatment: F (4,56) = 2.126, p = 0.0896, ANOVA). Analysis of effect size was not significant (Figure 4E). The effect of acidic saline on grip strength values persists when grip strength is normalized to total body weight, indicating that the effect is due to acidic saline treatment and not inherent group differences (Supplemental Figure 2A, treatment: F (1,14) = 8.089, p = 0.0130, ANOVA). Additionally, the effect of acidic saline treatment on grip strength does not persist through three weeks of assessment (Supplemental Figure 2C). This assay does not differentiate between ipsilateral and contralateral paws, which may be masking effects limited to a single side.

We used the incapacitance assay to test for changes in weight bearing at rest (Figure 4E). Although there was a significant effect of time on weight bearing (Figure 4F, time: F (2.664, 37.31) = 6.053, p = 0.0026, ANOVA), there was no effect of treatment or interaction. Analysis of effect size showed that there was a significant difference in weight bearing behavior between groups (Figure 4G: F(3,28) = 4.108, p = 0.0170, ANOVA). Although both groups favored the ipsilateral side (Figure 4 G-I), there was a shift in the weight bearing behavior of mice injected with acidic saline such that the preference for the ipsilateral side increased over time, whereas control mice displayed the opposite trend (Supplemental Figures 2E).

Finally, we used the rotarod assay to test for changes in endurance and muscle coordination. There was no effect of acidic saline injection on either of these measures (Supplemental Figure 2F).

### 3.4 Correlations between behavioral assays

#### 3.4.1. Comparisons to mechanical sensitivity

The acidic saline-induced model of chronic musculoskeletal pain has been robustly characterized using mechanical sensitivity measures (von Frey testing-cutaneous hyperalgesia, Tweezer-muscle hyperalgesia) (9, 11). All correlational data between each behavioral assay and mechanical sensitivity of the paw in the preclinical model of CWP can be found in Table 1. In summary, mechanical sensitivity of the paw (data from Merriwether et al., 2020; Figure 3 (11)) was robustly correlated with increased facial grimacing (Pearson r = 0.8367, p < 0.0001, R = 0.7000) and moderately correlated with preference for ipsilateral weight bearing following the first (Pearson r = -0.6134, p = 0.0155, R = 0.3763) but not second saline injection. Thermal sensitivity, grip strength, and coordination were not correlated with mechanical sensitivity of the paw.

**Table 1.** Correlations between mechanical sensitivity and pain and functional assays. Datasets were analyzed using Pearson r and two-tailed p-values. Effect sizes reported for each assay were used to calculate correlations. All significant correlations are bolded. CI = confidence interval.

| Outcome Domain | Behavioral Test | n | Pearson's correlation |  |  |  | Figure |
| --- | --- | --- | --- | --- | --- | --- | --- |
|  |  |  | r | 95% CI | p-value | R<sup>2</sup> |  |
| Spontaneous Pain | Facial Grimacing | 16 | **0.8367** | **0.5826-0.9418** | **<0.0001** | **0.7** | Fig 2. |
| Muscle Function | Grip Strength | 16 | 0.206 | -0.3227-0.6366 | 0.4442 | 0.0424 | Fig. 3 |
|  | **Weight Bearing – ipsilateral preference (Days 1-3)** | **16** | **-0.6134** | **-0.8505—0.1692** | **0.0115** | **0.3763** |  |
|  | Weight Bearing – ipsilateral preference (Days 6-9) | 16 | -0.1329 | -0.5898-0.3884 | 0.6237 | 0.0176 |  |
|  | Weight Bearing – contralateral preference (Days 1-3) | 16 | 0.2734 | -0.2571-0.6773 | 0.3055 | 0.0747 |  |
|  | Weight Bearing – contralateral preference (Days 6-9) | 16 | -0.0575 | -0.3227-0.6366 | 0.8324 | 0.0033 |  |
|  | Rotarod | 16 | -0.221 | -0.6587-0.3284 | 0.4286 | 0.0575 |  |
| Evoked Pain | Thermal Sensitivity | 16 | 0.288 | -0.2424-0.6857 | 0.2796 | 0.0829 | Fig. 3 |

#### 3.4.2. Inter-assay correlations

In the preclinical model of CWP, increased mechanical sensitivity of the paw following acidic saline injection was significantly correlated with increased facial grimacing (MGS post-movement: Pearson r = 0.8866, p < 0.0001, R = 0.7890; MGS Percent Change: r = 0.5381, p = 0.0212, R = 0.2896) after short term activity, indicating that mechanical hypersensitivity of the paw is strongly associated with spontaneous pain directly following activity.

Although not related to pain-behaviors, there were significant correlations between the grip strength and weightbearing (incapacitance) assays for muscle function tests. Decreases in grip strength were strongly associated with shifts in preference in weight bearing preference away from the ipsilateral side (Pearson r = 0.597, p = 0.015) and toward the contralateral side (Pearson r = -0.544, p = 0.029). Performance on the rotarod was not significantly associated with any other measures. Inter-assay correlations can be found in Table 2.

**Table 2.**
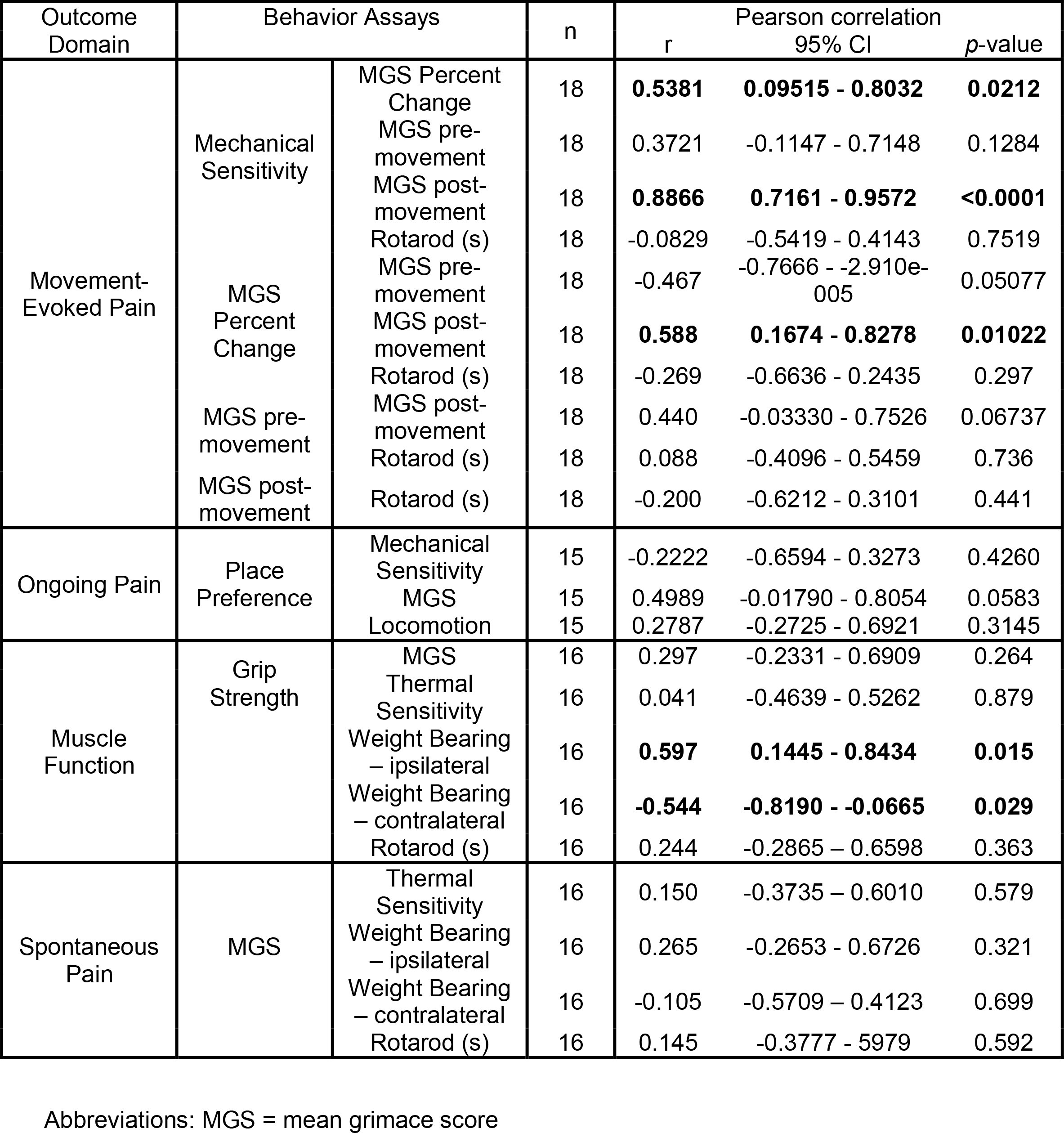
Correlations between behavioral assays for pain and muscle function. Datasets were analyzed via Pearson’s r correlations with two-tailed p-values.

### 3.5 Patient-Reported Outcomes

#### 3.5.1. Participant Characteristics

There were 289 women with fibromyalgia that met criteria for study inclusion and completed health questionnaires and pain tests and measures. The demographic data are presented in Table 3, with more detailed information about racial/ethnic identity, marital status, socioeconomic status, education level, and other variables of the clinical trial participants described elsewhere (17).

**Table 3.** Demographics and Clinical Variables. Data are presented as Mean (SE) and 95% confidence intervals. Spontaneous (Resting) pain intensity was measured using the Numeric Rating Scale (0-10); Movement-Evoked Pain was measured using the Numeric Rating Scale (0-10) during the Six-Minute Walk Test (6MWT) and Five Time Sit-to-Stand Test (5TSTS); Pain Interference and Pain Behavior were measured using PROMIS Short Forms _ and _; Physical activity (steps per day) was measured using accelerometry; The Six-Minute Walk Test (6MWT) physical function measure. Pain sensitivity was measured using the Pressure Pain Threshold (PPT) at the cervical, lumbar, and lower leg regions; Conditioned Pain Modulation (CPM) was measured using cold water immersion as the conditioning stimulus and PPT as the test stimulus. The value is the ratio of PPT values pre- and post-immersion.

| **Mean (SD)**<br>**Range** | **FM**<br>**N = 289** |
| --- | --- |
| Age, years | 48.21 (12.31)<br>20.20 – 69.20 |
| BMI (kg/m<sup>2</sup>) | 34.53 (8.76)<br>17.71 – 84.14 |
| Spontaneous (Resting) pain (0-10) | 6.18 (1.48)<br>4.00 – 10.00 |
| Movement-Evoked Pain – 6MWT (0-10) | 6.44 (1.84)<br>1.00 – 10.00 |
| Movement-Evoked Pain – 5TSTS (0-10) | 5.66 (2.21)<br>0.00 – 10.00 |
| PROMIS Pain Interference (T-score) | 64.81 (5.11)<br>52.50 – 78.30 |
| PROMIS Pain Behavior (T-score) | 60.64 (3.07)<br>52.10 – 68.30 |
| Physical Activity (steps per day) | 8256 (2906)<br>2024 – 18876 |
| 6MWT distance (feet) | 1340 (309)<br>300 - 2050 |
| 5TSTS (secs) | 14.14 (6.13)<br>5.50 – 53.00 |
| Cervical PPT (kPA) | 154.24 (71.41)<br>17.75 – 454.25 |
| Lumbar PPT (kPA) | 201.95 (100.69)<br>12.00 – 497.00 |
| Lower Leg PPT (kPA) | 183.24 (105.47)<br>25.00 – 666.50 |
| CPM Lumbar (Ratio) | 1.28 (0.42)<br>0.60 – 3.20 |
| CPM Lower Leg (Ratio) | 1.28 (0.46)<br>0.60 – 4.20 |

#### 3.5.2. Correlation Between Mechanical Pain Threshold (PPT) and Patient-Reported Outcomes in Women with FM

MEP measured during the 6MWT or 5TSTS test was associated with pain sensitivity at all body regions. Lower PPTs at the lumbar (⍰ =-0.193, p = 0.001) and right lower leg (⍰=-0.150, p = 0.011) were weakly associated with greater MEP measured during the 6MWT. Similarly, lower PPTs at the lumbar (⍰=-0.224, p ≤ 0.01) and right lower leg (⍰=-0.150, p = 0.005) were weakly to moderately associated with greater MEP measured during the 5TSTS test. Also, lower PPTs at the cervical site was moderately associated with greater MEP during the 5TSTS test (⍰=-0.175, p = 0.003). There was no significant association between cervical PPT and MEP during the 6MWT.

Pain sensitivity measured at the cervical site had inverse relationships with physical function and descending pain inhibitory control (i.e., CPM). Lower PPTs at the cervical site were weakly associated with 6MWT distance (⍰=-0.124, p = 0.036), 5TSTS test duration (⍰=0.143, p = 0.016), and less pain inhibition at the right lower leg site (CPM Leg; ⍰=-0.136, p = 0.032). The relationships between mechanical PPTs and resting (spontaneous) pain, pain interference, pain behaviors, physical activity levels (accelerometry), or pain inhibition at the lumbar or leg sites did not reach statistical significance (Table 4).

**Table 4.**
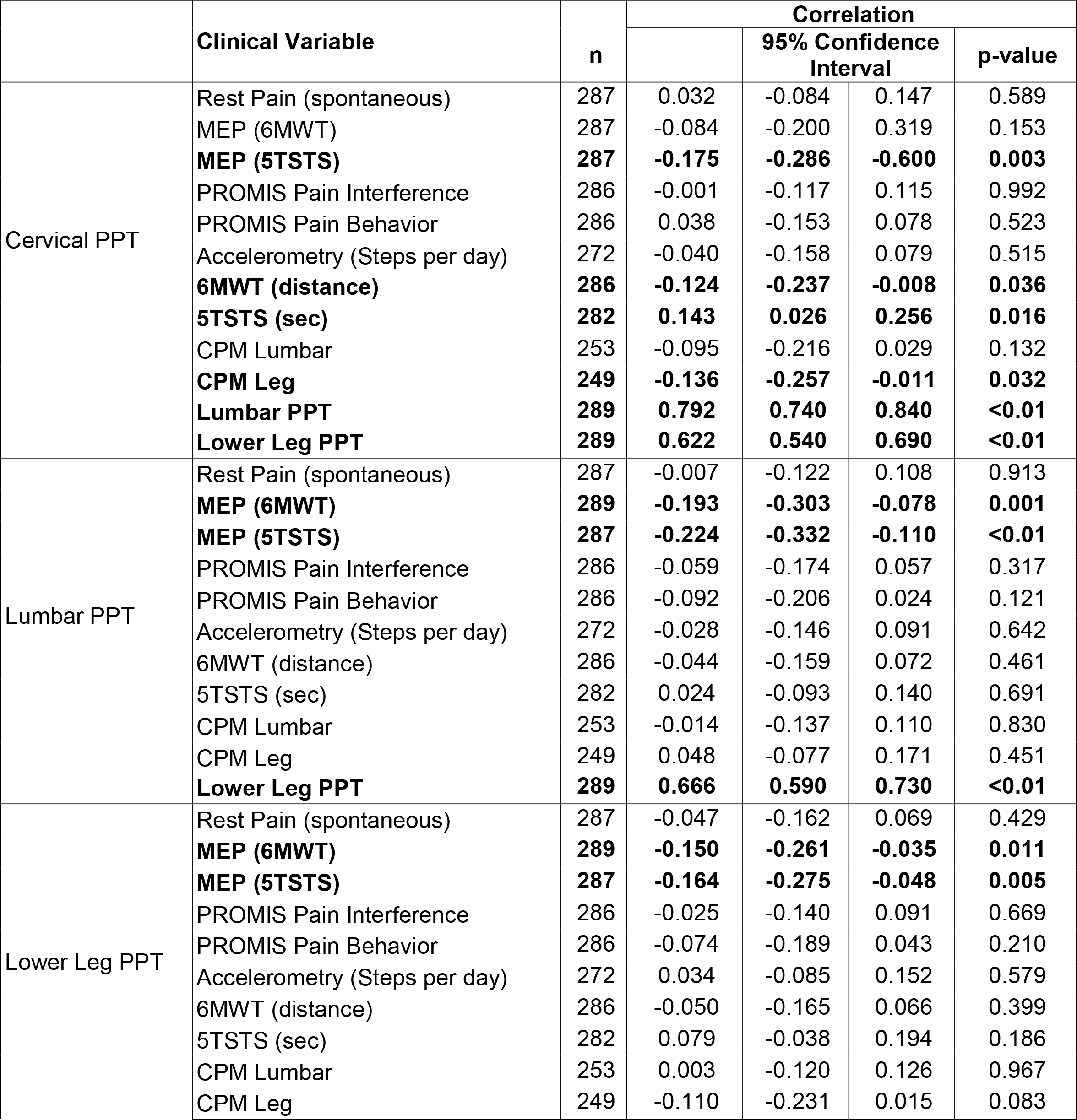
Correlations between mechanical pain threshold (PPT) and patient-reported outcomes **Abbreviations:** NRS Pain (rest), 6MWT, 5TSTS: Pain rating using the Numeric Rating Scale for pain at rest, during the six-minute walk test, and during the five time sit-to-stand test; BPI: Brief Pain Inventory, Severity and Interference subscales; PPT Cervical, Lumbar: Average measurements of pressure pain threshold (kPA) at the cervical and lumbar sites; CPM: Conditioned pain modulation (% change in PPT score) at the cervical and lumbar sites.

|  | Clinical Variable | n | Correlation |  |  |  |
| --- | --- | --- | --- | --- | --- | --- |
|  |  |  |  | 95% Confidence Interval |  | p-value |
| Cervical PPT | Rest Pain (spontaneous) | 287 | 0.032 | -0.084 | 0.147 | 0.589 |
|  | MEP (6MWT) | 287 | -0.084 | -0.200 | 0.319 | 0.153 |
|  | **MEP (5TSTS)** | **287** | **-0.175** | **-0.286** | **-0.600** | **0.003** |
|  | PROMIS Pain Interference | 286 | -0.001 | -0.117 | 0.115 | 0.992 |
|  | PROMIS Pain Behavior | 286 | 0.038 | -0.153 | 0.078 | 0.523 |
|  | Accelerometry (Steps per day) | 272 | -0.040 | -0.158 | 0.079 | 0.515 |
|  | **6MWT (distance)** | **286** | **-0.124** | **-0.237** | **-0.008** | **0.036** |
|  | **5TSTS (sec)** | **282** | **0.143** | **0.026** | **0.256** | **0.016** |
|  | CPM Lumbar | 253 | -0.095 | -0.216 | 0.029 | 0.132 |
|  | **CPM Leg** | **249** | **-0.136** | **-0.257** | **-0.011** | **0.032** |
|  | **Lumbar PPT** | **289** | **0.792** | **0.740** | **0.840** | **<0.01** |
|  | **Lower Leg PPT** | **289** | **0.622** | **0.540** | **0.690** | **<0.01** |
| Lumbar PPT | Rest Pain (spontaneous) | 287 | -0.007 | -0.122 | 0.108 | 0.913 |
|  | **MEP (6MWT)** | **289** | **-0.193** | **-0.303** | **-0.078** | **0.001** |
|  | **MEP (5TSTS)** | **287** | **-0.224** | **-0.332** | **-0.110** | **<0.01** |
|  | PROMIS Pain Interference | 286 | -0.059 | -0.174 | 0.057 | 0.317 |
|  | PROMIS Pain Behavior | 286 | -0.092 | -0.206 | 0.024 | 0.121 |
|  | Accelerometry (Steps per day) | 272 | -0.028 | -0.146 | 0.091 | 0.642 |
|  | 6MWT (distance) | 286 | -0.044 | -0.159 | 0.072 | 0.461 |
|  | 5TSTS (sec) | 282 | 0.024 | -0.093 | 0.140 | 0.691 |
|  | CPM Lumbar | 253 | -0.014 | -0.137 | 0.110 | 0.830 |
|  | CPM Leg | 249 | 0.048 | -0.077 | 0.171 | 0.451 |
|  | **Lower Leg PPT** | **289** | **0.666** | **0.590** | **0.730** | **<0.01** |
| Lower Leg PPT | Rest Pain (spontaneous) | 287 | -0.047 | -0.162 | 0.069 | 0.429 |
|  | **MEP (6MWT)** | **289** | **-0.150** | **-0.261** | **-0.035** | **0.011** |
|  | **MEP (5TSTS)** | **287** | **-0.164** | **-0.275** | **-0.048** | **0.005** |
|  | PROMIS Pain Interference | 286 | -0.025 | -0.140 | 0.091 | 0.669 |
|  | PROMIS Pain Behavior | 286 | -0.074 | -0.189 | 0.043 | 0.210 |
|  | Accelerometry (Steps per day) | 272 | 0.034 | -0.085 | 0.152 | 0.579 |
|  | 6MWT (distance) | 286 | -0.050 | -0.165 | 0.066 | 0.399 |
|  | 5TSTS (sec) | 282 | 0.079 | -0.038 | 0.194 | 0.186 |
|  | CPM Lumbar | 253 | 0.003 | -0.120 | 0.126 | 0.967 |
|  | CPM Leg | 249 | -0.110 | -0.231 | 0.015 | 0.083 |

#### 3.5.3. Correlation Between Pain and Other Patient-Reported Outcomes in Women with FM

Spontaneous (resting) pain. Spontaneous pain, measured as resting pain on a 11-point numeric rating scale, was associated with pain interference, pain behavior, and with objective measures of physical function. Spontaneous pain was moderately associated with more pain interference (⍰= 0.275, p<0.001) and more self-reported pain behaviors (⍰= 0.227, p<0.001). Greater spontaneous pain was associated with less distance walked during the 6MWT (⍰= -0.241, p<0.001) and longer times to complete the 5TSTS test (⍰= 0.160, p = 0.007).

MEP. MEP measured during the 6MWT and 5TSTS were correlated with physical activity level and with descending inhibitory pain control. Greater MEP measured during the 6MWT was negatively associated with CPM at the right lower leg site (⍰= -0.124, p < 0.001). Greater MEP measured during the 5TSTS test was associated with more steps per day (⍰= 0.170, p = 0.005) and was negatively correlated with the CPM at the lumbar site (⍰= -0.156, p = 0.013).

Pain Interference and Pain Behavior. Pain interference and pain behaviors were weakly to moderately associated with physical activity and performance-based functional measures. Greater pain interference was associated with less steps per day (⍰= -0.301, p < 0.001), less distance walked during the 6MWT (⍰= -0.175, p = 0.003), and a longer time to perform the 5TSTS (⍰= 0.133, p = 0.026). More pain behavior was associated with less distance walked during the 6MWT (⍰= -0.147, p = 0.013), and more time to complete the 5TSTS test (⍰= 0.206, p<0.001).

Descending Pain Inhibition (CPM). Less CPM, which is indicative of impaired descending inhibitory pain control, was associated with longer times to perform the 5TSTS test (⍰= -0.132, p= 0.039). There were no significant correlations between CPM, spontaneous pain, pain interference, pain behavior, or level of physical activity. Correlations between pain and other patient-reported outcomes can be found in Table 5.

**Table 5.** Correlation of Patient-Reported Outcomes with Pain **Abbreviations:** NRS Pain (rest), 6MWT, 5TSTS: Pain rating using the Numeric Rating Scale for pain at rest, during the six-minute walk test, and during the five time sit-to-stand test; BPI: Brief Pain Inventory, Severity and Interference subscales; PPT Cervical, Lumbar: Average measurements of pressure pain threshold (kPA) at the cervical and lumbar sites; CPM: Conditioned pain modulation (% change in PPT score) at the cervical and lumbar sites.

| Pain variable | Clinical variable | n | Pearson correlation |  |  |  |
| --- | --- | --- | --- | --- | --- | --- |
|  |  |  |  | 95% Confidence Limits |  | p-value |
| Rest Pain<br>(spontaneous) | MEP (6MWT) | 287 | 0.034 | -0.082 | 0.149 | 0.567 |
|  | MEP (5TSTS) | 285 | 0.028 | -0.088 | 0.143 | 0.632 |
|  | **PROMIS Pain Interference** | **288** | **0.275** | **0.016** | **0.380** | **<0.001** |
|  | **PROMIS Pain Behavior** | **288** | **0.227** | **0.113** | **0.335** | **<0.001** |
|  | Accelerometry (Steps per day) | 274 | -0.074 | -0.190 | 0.050 | 0.221 |
|  | **6MWT (distance)** | **288** | **-0.241** | **-0.348** | **-0.127** | **<0.001** |
|  | **5TSTS (sec)** | **284** | **0.160** | **0.040** | **0.272** | **0.007** |
|  | CPM Lumbar | 251 | -0.100 | -0.221 | 0.024 | 0.114 |
|  | CPM Lower Leg | 247 | -0.057 | -0.180 | 0.068 | 0.372 |
| MEP (6MWT) | **MEP (5TSTS)** | **287** | **0.536** | **0.442** | **0.618** | **<0.001** |
|  | PROMIS Pain Interference | 286 | -0.045 | -0.162 | 0.071 | 0.447 |
|  | PROMIS Pain Behavior | 286 | -0.050 | -0.165 | 0.066 | 0.400 |
|  | Accelerometry (Steps per day) | 272 | 0.075 | -0.045 | 0.192 | 0.220 |
|  | 6MWT (distance) | 286 | -0.067 | -0.182 | 0.049 | 0.261 |
|  | 5TSTS (sec) | 282 | 0.078 | -0.040 | 0.193 | 0.189 |
|  | CPM Lumbar | 253 | -0.106 | -0.226 | 0.019 | 0.093 |
|  | **CPM Lower Leg** | **249** | **-0.124** | **-0.245** | **0.001** | **0.051** |
| MEP (5TSTS) | PROMIS Pain Interference | 284 | -0.005 | -0.121 | 0.111 | 0.932 |
|  | PROMIS Pain Behavior | 284 | 0.035 | -0.082 | 0.151 | 0.555 |
|  | **Accelerometry (Steps per day)** | **271** | **0.170** | **0.051** | **0.284** | **0.005** |
|  | 6MWT (distance) | 284 | 0.029 | -0.088 | 0.145 | 0.624 |
|  | 5TSTS (sec) | 280 | 0.003 | -0.114 | 0.120 | 0.954 |
|  | **CPM Lumbar** | **249** | **-0.159** | **-0.275** | **-0.032** | **0.003** |
|  | CPM Lower Leg | 245 | -0.111 | -0.231 | 0.016 | 0.084 |
| PROMIS Pain<br>Interference | **PROMIS Pain Behavior** | **288** | **0.621** | **0.537** | **0.693** | **<0.001** |
|  | **Accelerometry (Steps per day)** | **273** | **-0.301** | **-0.407** | **-0.186** | **<0.001** |
|  | **6MWT (distance)** | **287** | **-0.175** | **-0.286** | **-0.060** | **0.003** |
|  | **5TSTS (sec)** | **283** | **0.133** | **0.017** | **0.245** | **0.026** |
|  | CPM Lumbar | 251 | 0.016 | -0.108 | 0.140 | 0.802 |
|  | CPM Lower Leg | 247 | 0.028 | -0.097 | 0.152 | 0.665 |
| PROMIS Pain<br>Behavior | Accelerometry (Steps per day) | 273 | -0.084 | -0.201 | 0.035 | 0.164 |
|  | **6MWT (distance)** | **287** | **-0.147** | **-0.259** | **-0.031** | **0.013** |
|  | **5TSTS (sec)** | **283** | **0.206** | **0.090** | **0.316** | **<0.001** |
|  | CPM Lumbar | 251 | -0.042 | -0.165 | 0.082 | 0.504 |
|  | CPM Lower Leg | 247 | 0.006 | -0.119 | 0.131 | 0.926 |
| CPM Lumbar<br>(%) | Accelerometry (Steps per day) | 236 | -0.052 | -0.179 | 0.076 | 0.428 |
|  | 6MWT (distance) | 251 | 0.062 | -0.062 | 0.185 | 0.327 |
|  | 5TSTS (sec) | 248 | -0.040 | -0.164 | 0.085 | 0.531 |
|  | **CPM Lower Leg** | **249** | **0.533** | **0.431** | **0.622** | **<0.001** |
| CPM Leg (%) | Accelerometry (Steps per day) | 232 | 0.008 | -0.121 | 0.137 | 0.898 |
|  | 6MWT (distance) | 247 | 0.117 | -0.008 | 0.239 | 0.067 |
|  | **5TSTS (sec)** | **244** | **-0.132** | **-0.254** | **-0.006** | **0.039** |

## 4. Discussion

The primary aim of the current study was to validate a range of behavioral assessments in a preclinical model of FM and compare them to clinical outcomes from the Fibromyalgia Activity Study with TENS (FAST) clinical trial and examine relationships between select patient-reported and measured outcomes for women with FM (17). The current study shows that in the preclinical model of CWP, female mice exhibit a range of pain behaviors: movement-evoked mechanical sensitivity and facial grimacing, spontaneous pain, decreased muscle strength, and altered weight bearing behavior. There were not lasting changes in thermal sensitivity or muscle performance. Similarly, women with fibromyalgia reported resting and movement-evoked pain, showed decreases in pain threshold, decreased muscle performance and reduced activity levels. Mechanical sensitivity of the paw, which has been previously characterized in this model (9, 11, 12) was strongly associated with facial grimacing behavior, ipsilateral weight-bearing preference, and movement-evoked pain. In women with fibromyalgia, we found that mechanical pain thresholds, measured via PPT, were moderately correlated with MEP and performance-based physical function. Notably, resting pain was a moderate predictor of both pain interference and pain behaviors, whereas MEP was not. Furthermore, in the animal model we found that although behavioral tests for muscle performance did not correlate with pain behaviors, there were inter-assay correlations for changes in muscle function such that decreases in grip strength were strongly associated with prolonged shifts in weight bearing preference for the side of injection. In contrast, women with FM showed strong correlations between pain interference and PROMIS pain behavior and physical activity measures. This study adds to the FM literature by providing a battery of pain behavioral assessments in a preclinical model that provides a framework for the selection of pain assays that could foster more effective better translation of preclinical findings when testing novel therapeutics, and potentially have increased translational value based on the assessments performed in clinical settings.

Mechanical pain thresholds have weak to moderate correlations with MEP in preclinical and clinical models of CWP and FM. Patients with FM often report that therapeutics do not effectively alleviate MEP and that it is a major limiting factor in patient participation in exercise-based interventions (38, 39) which have been shown to be effective at both the preclinical and clinical stages (17, 19, 40, 41); However, MEP is not well defined, is not consistently measured in clinical settings (42, 43), and disproportionately burdens older adults and select racial and ethnic groups (44–46). Preclinical settings rarely measure MEP, as such there is no standardized methodology for assessing this type of pain. We developed a novel paradigm for assessing MEP in mice by measuring mechanical sensitivity of the paw and facial grimacing before and after activity in mice with CWP. A similar paradigm was used in Khataei et al. (2020) in which exhaustive exercise on a treadmill induced a decrease in muscle withdrawal thresholds (41); However, this study was performed on naïve mice and not on mice with CWP. Other studies using this model of CWP have assessed the analgesic effects of low intensity exercise on mechanical sensitivity, but pain assessments did not occur immediately following exercise and these assessments were made over several weeks (40, 47). Our study provides evidence that MEP can be successfully measured in a model of CWP using facial grimacing; However, future studies should assess additional measures of pain following activity such as muscle withdrawal thresholds and thermal sensitivity.

Using our novel approach, the preclinical model shows robust MEP, thus, the efficacy of therapeutics in mitigating MEP could be effectively determined. Previous work has shown that transcutaneous electric nerve stimulation (TENS) successfully reduces MEP in women with FM (17), and reduced pain in the preclinical model of FM (8). Although mice with CWP had decreased bilateral sensitivity, this was not changed following movement assessments. One of the limitations of testing mechanical withdrawal thresholds in rodents is that chronic pain models often see a basement effect, in which additional decreases in withdrawal thresholds cannot be detected due to the limited range of the test. Using facial grimacing, we were able to see an increase in spontaneous pain, as the animals had not reached their maximum prior to the assessment of MEP. Although there were correlations between mechanical sensitivity of the paw and facial grimacing during MEP assessment, use of facial grimacing allows for increased sensitivity to detect changes in MEP due to basement effects observed in mechanical withdrawal thresholds. Testing interventions to reduce MEP at the preclinical stage has increased translational value and would potentially increase the efficacy and success rates of preclinical trials, particularly when using assessments of spontaneous MEP in addition to mechanical sensitivity.

Pain is an inherently subjective and multifaceted experience, defined by both sensory nociception and negative emotional valence that accompanies it (48). Patients with chronic pain often adjust their expectations and define for themselves a new “normal” level of pain, thus reducing the negative emotional state that typically accompanies nociception (49, 50). Patients do not experience a constant level of pain, rather they tend to experience flare-ups in pain that are typically triggered by outside sources such as stress, activity, and emotional state (51). This is fundamentally different from the pain behaviors measured in animal models, where researchers are less able to measure the quality and impact of pain and focus primarily on nociception (2, 3). The use of non-evoked pain measures in preclinical models helps to bridge this gap, as spontaneous pain measures are more closely related to the subjective nature of pain than measures of nociception (37, 52). Facial grimacing has been used to assess affective components of pain across multiple pain models, including migraine and post-surgical pain, and is often used to assess the efficacy of analgesics (53–56). Even in studies of spontaneous pain using animal models, interpretation of rodent behaviors has to be considered in the context of prey behavior, as mice are prone to adjusting their behavior for survival (37). Typically, facial grimacing in mice occurs during the acute phase of chronic pain and does not last more than a few days (37, 57); however, in the acid-induced model of FM we found that facial grimacing persists three weeks beyond the induction of CWP. The use of affective measures of pain in the CWP model may provide a more direct measure of fluctuations in resting and affective pain states compared to evoked measures alone.

Physical function deficits in women with fibromyalgia have been reported across several studies, with pain and fatigue representing major barriers to participation in physical activities (58, 59). Relationships between pain and physical performance in individuals with FM are heterogeneous, influenced by perceived function and fatigue (13, 14); However, the patient population represented in our study showed associations between resting pain and deficits in physical function measured by the 6MWT and 5TSTS. In the animal model, there was not an association between spontaneous (resting) pain measured via facial grimacing and physical function measures (grip strength and weight bearing). There were, however, correlations between mechanical sensitivity of the paw and weight bearing behaviors. Similarly, individuals with FM showed associations between PPT measures and physical function deficits.

Our intent in this study was to assess the translational validity of pain behaviors in a preclinical model of CWP. The assays performed in the preclinical portion of this study were all conducted in a testing room and examined pain behaviors in various contexts such as evoked pain, spontaneous pain, and muscle strength. We acknowledge that animal behavior in this context may not completely reflect home-cage behavior, such as voluntary wheel-running, cage-lid hanging, and general home-cage activity, which reduces stress from test that occur out of the animal’s home environment (60, 61). Home-cage behaviors may also be more translational to quality-of-life assessments. This study also exclusively focused on female animals, as the patient population from the FAST study was conducted on female patients with FM. We acknowledge that although FM presents predominately in females, including males in both preclinical and clinical studies would improve generalizability of these findings and potentially highlight sex differences in the pathology of FM. Furthermore, patients with FM often have comorbid conditions, such as osteoarthritis, rheumatoid arthritis, and migraine, that may influence pain behaviors in patients and introduce variability (13, 14, 17, 51).

In conclusion, the acidic saline model of FM successfully recapitulates many of the symptoms experienced by patients with FM, with similar relationships between symptom subsets. Future studies on preclinical FM models should consider expanding the battery of behavioral assays when testing the efficacy of therapeutics. Therapies that reduce MEP in particular are of high value, as reductions in MEP may improve patient outcomes in exercise-based treatments. Preclinically, assessment of MEP should focus primarily on spontaneous pain rather than relying solely on changes in mechanical sensitivity to detect changes in pain states.

## Acknowledgements

Animal work was supported by NIH grant K22NS096030 (MDB), American Pain Society Future Leaders Grant (MDB), Rita Allen Foundation Award in Pain (MDB) and The University of Texas System Rising STARS program research support grant (MDB). This work is supported by NIH UM1 AR06338 and NIH UM1 AR06338-S1. Human study data was collected and managed using REDCap electronic data capture tools hosted at the University of Iowa (supported by NIH 54TR001013). FAST data collection was completed at the CTSA at University of Iowa (supported by NIH U54TR001356) and Vanderbilt University (supported by NIH UL1TR000445). The authors would like to acknowledge Jessica Danielson for her contributions to data collection for the human study. The authors would also like to thank Michelle Vo and Gabrielle Cox for their technical assistance, as well as current and past members of the Neuroimmunology and Behavior Lab. Graphics were created using BioRender.com.

---

**Supplemental Figure 1. Pain behaviors tested through three weeks following acidic saline injections.** (A) Facial grimacing was tested for three weeks after the first saline injection. Facial grimacing is significantly increased for three weeks compared to controls (time x treatment: F (8, 48) = 3.318, p = 0.0043, Repeated Measures Two-Way ANOVA, n=8/group through Day 9 and n=4/group through Day 21). (B) Effect size for facial grimacing for days 12-21 is significantly increased compared to controls (t (6) = 7.667, p = 0.0003, unpaired t-test, two-tailed, n=4/group). (C) Thermal sensitivity is decreased three weeks following the first saline injection (time x treatment: F (8,80) = 4.624, p = 0.0001, Repeated Measures Two-Way ANOVA, n=8/group through Day 9 and n=4/group through Day 21). (D) Effect size analysis for thermal sensitivity for days 12-21 (n=4/group). Black arrowheads indicate saline injections. MGS = mean grimace score. *p<0.05, **p<0.01, ***p<0.001, ****p<0.0001.

**Supplemental Figure 2. Muscle performance behaviors tested through three weeks following acidic saline injections.** (A) Grip strength normalized to individual mouse weights (treatment: F (1,14) = 8.089, p = 0.0130, Repeated Measures Two-Way ANOVA, n=8/group). (B) Effect size analysis for normalized grip strength. (C) Grip strength for days 12-21 (n=4/group). (D) Effect size for grip strength from days 12- 21. (E) Weight bearing behaviors for Days 15-21 (n=4/group). (F) Time spent on the rotarod per trial at baseline, Day 0 after the first injection, and Day 10 after the development of CWP (n=8/group). Black arrowheads indicate saline injections. **p<0.01.

