## Supplementary figures and images for "Translating outcomes from the clinical setting to preclinical models: chronic pain and functionality in chronic musculoskeletal pain"

### Supplemental Figures

Supplemental Figure 1

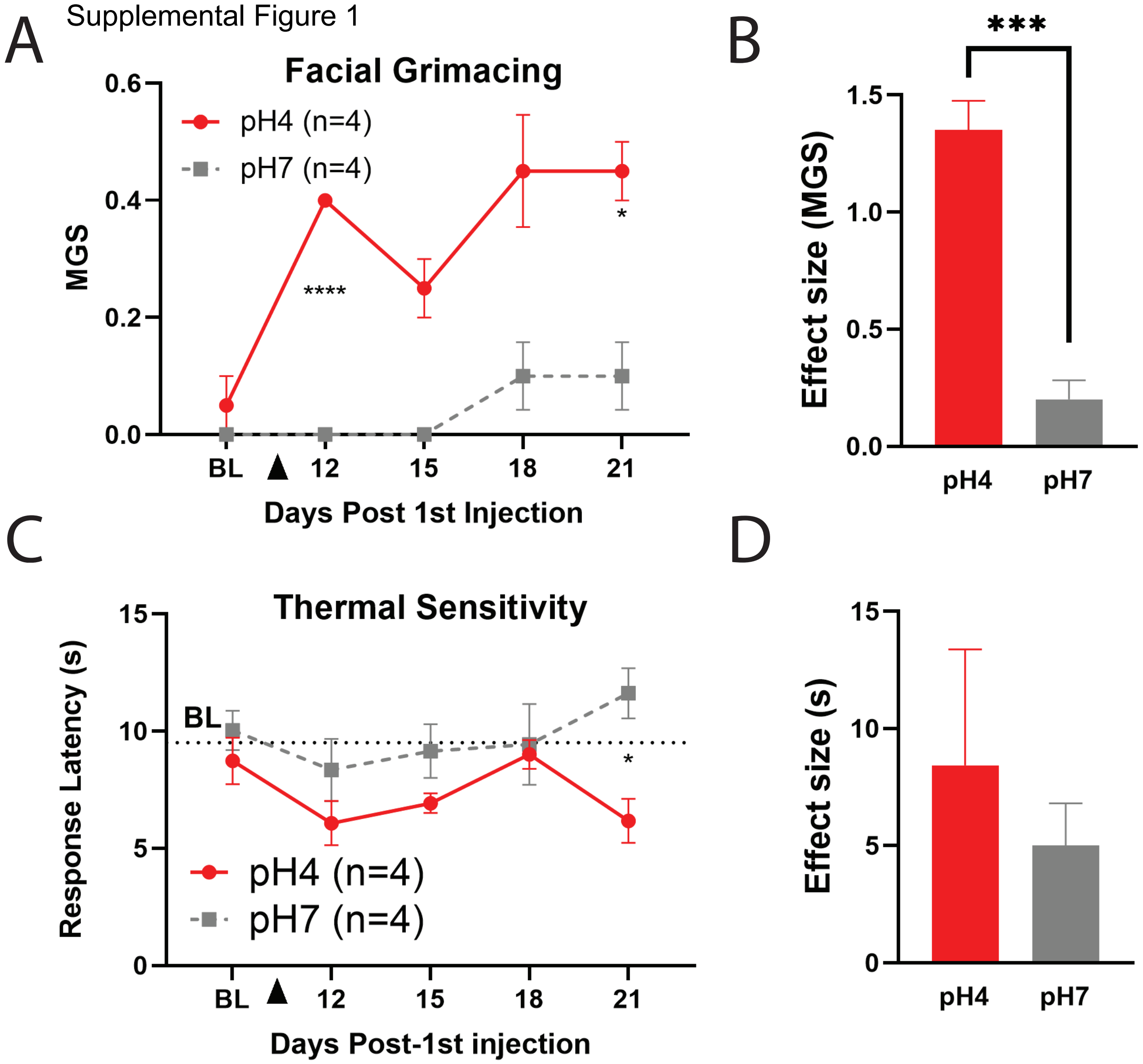

Supplemental Figure 2

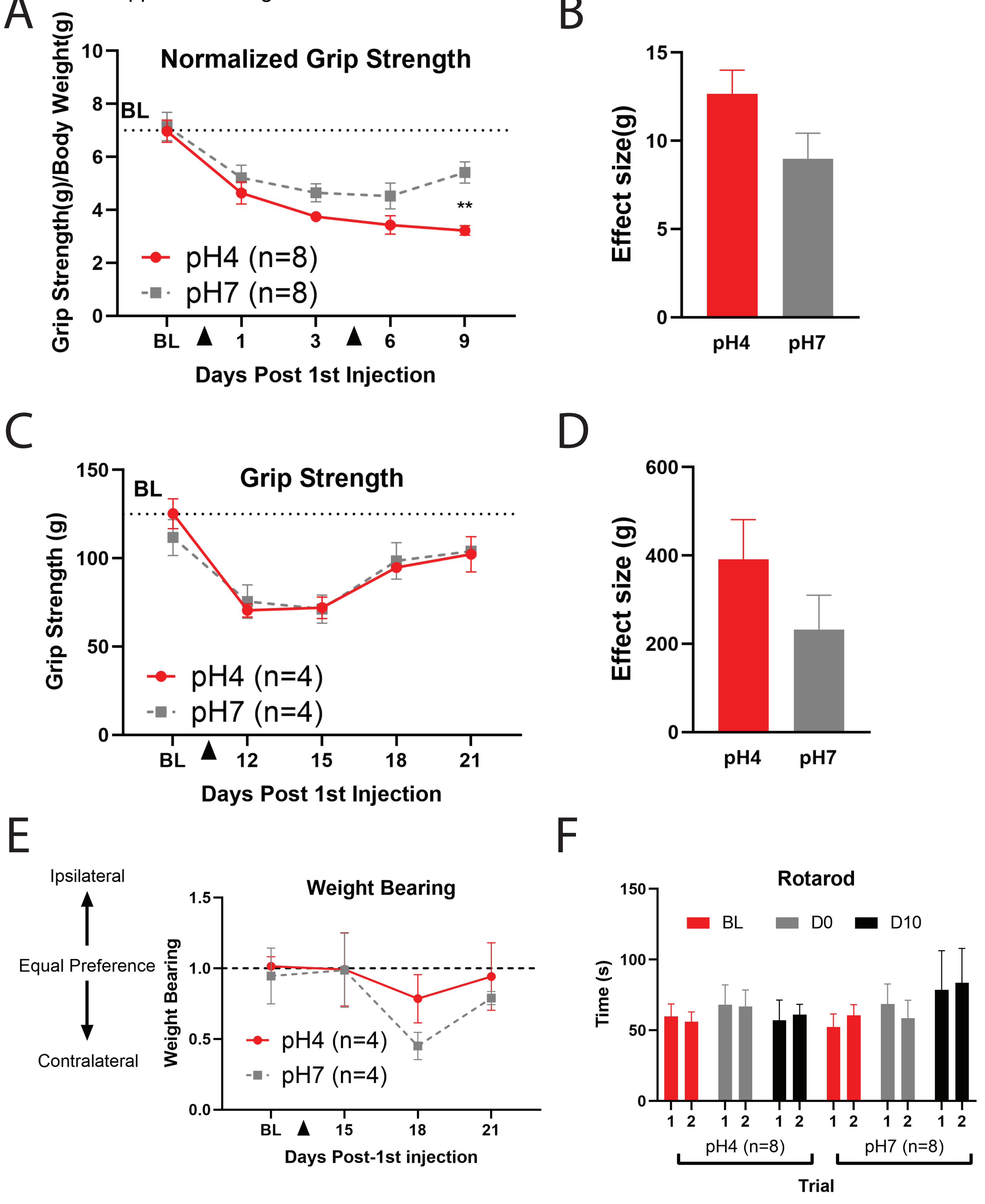
